# Modelling the interaction between Infectious Bursal Disease Virus VP3 polypeptide and dsRNA: identification of key residues for ribonucleoprotein assembly and virus replication

**DOI:** 10.1101/2020.08.06.240028

**Authors:** Idoia Busnadiego, Maria T Martín, Diego S Ferrero, María G Millán de la Blanca, Laura Broto, Elisabeth Díaz-Beneitez, Daniel Fuentes, Dolores Rodríguez, Nuria Verdaguer, Leonor Kremer, José F. Rodríguez

## Abstract

The interaction of the structural VP3 polypeptide of infectious bursal disease virus (IBDV) with virus-encoded dsRNA is essential both for the assembly of ribonucleoprotein complexes responsible for genome transcription and replication and for the evasion of host’s antiviral responses. Surface plasmon resonance analysis allowed us to determine the kinetic constants of the VP3-dsRNA interaction as well as to map the VP3 dsRNA bipartite dsRNA binding domain (dsRBD), uncovering the specific role of the previously described Patch1 and Patch2 dsRB subdomains. Here we show that the Patch1 domain plays a primary binding function while Patch2 exerts a subordinate role stabilizing VP3-dsRNA complexes. The use of a set of VP3 mutant versions facilitated the identification of K99 and K106 within Patch1 as the essential residues for the formation of VP3-dsRNA complexes. Furthermore, replacement of either one of these two residues by aspartic acid completely thwarts both evasion from host’s sensors and virus replication. Data presented here allow us to propose a VP3-dsRNA interaction model that should help to further elucidate the mechanics of IBDV morphogenesis and genome packaging as well as to better understand how VP3 counteracts recognition of virus-encoded dsRNA by specialized host’s sensors.

## INTRODUCTION

Double-stranded (ds) RNA duplexes are ubiquitously found in all live entities. dsRNA modules are indeed essential for a wide variety of biological processes, e.g. mRNA splicing and transport, gene transcription and mRNA translation. The multiple roles played by dsRNA modules entail their interaction with specific protein partners. Ribosomes and spliceosomes are good examples of the importance of these interactions for the maintenance of life. Paralleling their paramount biological significance, dsRNA-protein interactions have been analysed using a wide variety of experimental approaches, thus multiple dsRNA-binding proteins have been structurally characterized leading to a better understanding on how proteins recognize and interact with RNA duplexes (Saunders and Barber 2003; Wang et al. 2015).

Viruses, especially those harbouring dsRNA genomes, are utterly dependent upon dsRNA-protein interactions both to successfully completing their replication program and preventing detection by host-encoded dsRNA pattern recognition receptors (PRRs) and dsRNA-dependent antiviral effectors that constitute a major component of the cellular innate antiviral response (Jensen and Thomsen 2012).

The *Birnaviridae* family comprises a group of non-enveloped icosahedral viruses harbouring bi-segmented, double-stranded RNA (dsRNA) genomes (Delmas et al. 2004). Members of this family infect insects, aquatic fauna and birds. Within this family, Drosophila X (DXV), pancreatic necrosis (IPNV), blotched snakehead (BSV) and infectious bursal disease virus (IBDV) are prototype members of the entomo-, aqua-, blosna- and avibirnavirus genera, respectively (Delmas et al. 2004).

IBDV, the best characterized member of this family, infects domestic chickens (*Gallus gallus*) causing an acute immunosuppressive disease that imposes severe loses to the poultry industry worldwide (van den Berg et al. 2000). The IBDV major structural components namely VP2 (the capsid polypeptide) and VP3 (a multifunctional polypeptide), are encoded by the polyprotein open reading frame (Kibenge et al. 1997). The polyprotein undergoes a co-translational, self-proteolytic processing that releases pVP2 (the capsid polypeptide precursor) along with VP4 (the viral protease) and VP3 (Sánchez and Rodríguez 1999). VP3 is multifunctional participating in several processes during IBDV replication, i.e. acting as a scaffolding element during particle morphogenesis (Maraver et al. 2003; Oña et al. 2004; Saugar et al. 2010), activating the virus-encoded RNA-dependent RNA polymerase (RdRP, also known as VP1) (Garriga et al. 2007; Ferrero et al. 2015), and interacting with both dsRNA genome segments to form ribonucleoprotein (RNP) complexes that occupy the inner particle space (Luque et al. 2009).

Prototypical dsRNA viruses, e.g. members of the *Reoviridae* family, enclose their multipartite genomes into a conserved icosahedral structure, known as transcriptional core, which remains intact throughout the replication cycle. This structure holds the enzymatic machinery required for genome replication, transcription and mRNA extrusion whilst providing an efficient shelter against dsRNA host sensor proteins (Lawton et al. 1997). In a sharp structural and functional contrast, Birnaviruses lack replicative cores (Böttcher et al. 1997; Coulibaly et al. 2005; Castón et al. 2008). This structure is functionally replaced by RNP complexes built by the genome dsRNA segments associated to VP3 dimers. The third, and minor, RNP component is the virus-encoded RNA polymerase (VP1), which is found in two molecular forms, i.e. as a free polypeptide and as VPg, covalently linked to the 5’-ends of both genome dsRNA segments (Hjalmarsson et al. 1999; Luque et al. 2008). RNPs are transcriptionally active, *in vitro* and *ex vivo*, and act as transcriptional/replication devices capable of triggering a productive infection in the absence of the capsid protein (Luque et al. 2008; Dalton and Rodríguez 2014).

Like other dsRNA binding proteins, e.g. NS1 from influenza virus (Tan and Katze 1998) and E3 from vaccinia virus (VACV) (Romano et al. 1998), *in vitro* assays indicate that VP3 shields viral dsRNA from cellular dsRNA sensors, thus preventing the activation of at least two major components of the host’s innate antiviral protein arsenal, namely the protein kinase RNA-activated (PKR) (Busnadiego et al. 2012) and the melanoma differentiation-associated protein 5 (MDA5) (Ye et al. 2014). Additionally, we have also found that VP3 acts as an efficient anti-silencing protein in an experimental virus/plant model (Valli et al. 2012).

According to our preliminary mapping, the VP3 polypeptide holds a bipartite dsRNA binding domain (dsRBD) located at the central region of the protein (Valli et al. 2012). The dsRBD encompasses two surface exposed electropositive regions, termed Patch1 and Patch2, located at the same protein face. Each region holds four electropositive residues: Patch1 (K99, R102, K105 and K106) and Patch2 (R159, R168, H198 and R200) (Valli et al. 2012). The central VP3 region harbouring the dsRBD folds in two a-helical modules connected by a long and flexible hinge and is organized as a swapped dimer (Casañas et al. 2008).

So far, attempts to solving the crystal structure of VP3 bound to dsRNA have failed, thus hindering the possibility of obtaining precise information about the VP3-dsRNA interaction mechanism. Here, using surface plasmon resonance (SPR) analysis, we have characterized the interaction of wild-type and mutant versions of the VP3 polypeptide with synthetic RNA duplexes. Our results provide novel information about the VP3-dsRNA binding kinetics and show that the Patch1 region acts as the primary binding site while Pacth2 plays a secondary role stabilizing the VP3-dsRNA interaction. We have also identified Patch1 K99 and K106 as the critical residues for dsRNA binding as mutations affecting either one of them abrogate dsRNA binding as well as the capacity of the protein to shield dsRNA from cellular sensors and evade the host’s innate response. We also show that such mutations abolish IBDV infectivity. Based on the previously described VP3 dimer crystal structure, we propose a VP3-dsRNA complex model perfectly fitting the interaction data described here.

## RESULTS

### *In silico* VP3-dsRNA interaction model

As described above, our attempts to solve the crystal structure of the VP3-dsRNA complex have not yet been successful. However, the previously described atomic structure of VP3 (Casañas et al. 2008) together with our preliminary mapping of the VP3 dsRBD (Valli et al. 2012) prompted us to generate a hypothetical interaction model. Here, we propose an *in silico* model allowing a feasible electrostatic docking of the VP3 protein (pdb: 2r18) and the A-form of the dsRNA helix (pdb: 1RNA). As shown in Fig. 1A, the electronegative side chains of Patch1 residues K99 and K106, located on α-helix 2 of VP3, are spaced by 12.3 Å. This distance is rather close to the 13.9 Å gap spacing the OH^-^ groups from 5’-phosphates of nucleotides n1 and n9c (the latter corresponding to the complementary RNA strand) within the major groove of the dsRNA helix. This suggests that both phosphates could be efficiently trapped by K99 and K106 residues. In this context, the closely located R126 (VP3 α-helix 1) and the hydrophobic I103 residue (VP3 α-helix 2) could also play a significant role stabilizing the complex, either assisting K99/K106-mediated interactions and/or blocking the displacement of the dsRNA. The Patch2 subdomain, placed below Patch1 within the same face of the VP3 monomer, would also contribute to stabilizing dsRNA binding. Both dsRBD subdomains are off centered the vertical VP3 axis, with Patch2 facing the half of α-helix 2 harboring the seemingly crucial Patch1 K99 and K106 residues (Fig. 1B). The average distance between K99 and the clustered electropositive residues forming Patch2 is of 37.3 Å, fairly close to the 37.7 Å gap spacing the OH^-^ groups from the 5’-phosphates of nucleotides n1 and n17c at the major dsRNA helix groove.

**Figure 1.**
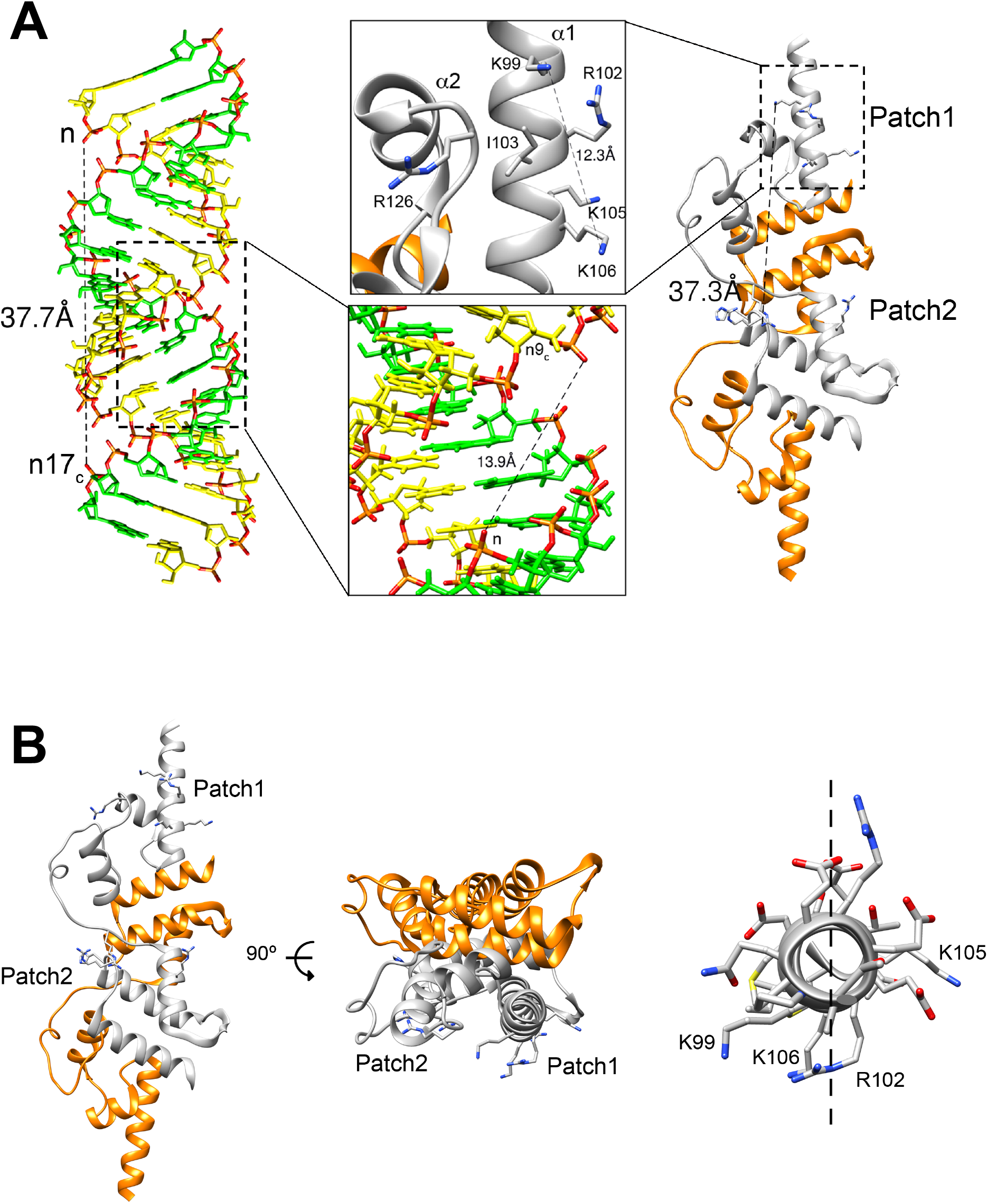
In silico VP3-dsRNA interaction model. (A) Matching phosphate distances in dsRNA and VP3 dsRB Patch1 and Patch2 subdomains. The left panel shows a stick representation of a dsRNA structure in a standard A-conformation (PDB id.6BU9). The bases and sugars of each chain are depicted in yellow and green, with the phosphate groups shown in orange and red for the phosphate and oxygen atoms, respectively. The right panel shows a ribbon representation of IBDV VP3, highlighting the positive residues forming Patch1 and Patch2. The distance between K99 in Patch1 and R159 in Patch 2 is 37.3 Å (dashed line), very close to the distance between phosphate groups in a 17 nucleotides dsRNA duplex (labelled in the left panel). The upper central panel shows a close-up of the VP3 Patch1 subdomain. The dashed-line indicates the distance between residues K99 and K106 that is almost coincident with the distance linking the phosphate groups of complementary chains in a 9-bp dsRNA region, as shown in the lower central panel. (B) **VP3 dsRBD subdomains.** Left panel, ribbon representation of IBDV VP3 dimer highlighting the electropositive dsRBD residues forming Patch1 and Patch2, and each VP3 chain in orange and grey, respectively. Central panel, top view of VP3. Right panel, top view of α-helix 1, the dashed line indicates the central part of the secondary structure with most of the positive residues facing Patch2.

According to the model, stable VP3-dsRNA complex formation would be strictly dependent upon the length of the interacting dsRNA helix providing a suitable platform for the correct positioning of both VP3 dsRBD subdomains. According to the model, a perfectly base-paired dsRNA helix formed by 17-bp should support the engagement of both Patch1 and Patch2 subdomains whilst a shorter one formed by 9-bp would allow a weaker interaction with Patch1.

Indeed, these theoretical helix sizes are prone to variations due to the inherent instability of the helix, which is especially high at both ends. This phenomenon, known as “breathing”, has been thoroughly described in the DNA helix and reflects spontaneous local conformational fluctuations within double-stranded DNA leading to the breaking of base pairs at temperatures below the melting temperature (Fei and Ha 2013). Indeed, the breathing of dsRNA analytes used in our study might impose some deviation of experimental data with respect to the proposed model.

### Kinetic analysis of the VP3-dsRNA interaction

Previous work from our laboratory showed that hVP3, a recombinant version of the VP3 polypeptide harbouring an N-terminal polyhistidine tag, efficiently binds purified IBDV dsRNA genomic segments (ca. 3 Kb) as well as short (21- and 26-bp) synthetic dsRNA duplexes (Valli et al. 2012). That study was based on the use of electrophoretic mobility assays (EMSA), an experimental approach based on equilibrium analysis that does not provide information about protein-dsRNA binding kinetics. Hence mechanistic details about the VP3-dsRNA interaction remained unknown.

Indeed, testing the interaction model described above required a more sophisticated approach allowing to compare the performance of different RNA duplexes and mutant VP3 versions. For this, we resorted to the use of SPR, a highly sensitive approach providing real time quantitative binding kinetics data and allowing the detection of weak protein-dsRNA interactions (Myszka 1997).

In order to obtain a comprehensive analysis of the hVP3-dsRNA interaction, biotinylated RNA duplexes of different lengths (10-, 12-, 14-, 16-, 18-, 20-, 22-, 24-, 32- and 40-bp) were captured on the surface of different flow cells of a streptavidin-coated sensor-chip. Analyses were then performed by injecting different concentrations (ranging from 0.313 to 20 nM) of affinity-purified hVP3 onto the RNA duplexes captured on sensor-chips following the protocol described in the Material and Methods section.

Results of this analysis are shown in table 1 and supplementary figure 1. Although our model predicts the binding of VP3 dimers to a helix comprising 10-bps, experimental data show that this is not the case, probably reflecting helix end’s instability and the consequent reduction of the available dsRNA length.

Complex formation was detected with all other tested RNA duplexes (Suppl. Fig. 1). As shown in the corresponding sensorgrams, responses during the hVP3 association and dissociation phases are fast and highly dependent on protein concentration. Recoded information was used to calculate kinetics data using a simple 1:1 Langmuir model including a term for mass transport deficiency. Noteworthy, as shown in Suppl. Fig. 1, data gathered with the 40-bp analyte do not fit the 1:1 Langmuir model, hence precluding the calculation of reliable kinetics data. Indeed, this observation strongly suggests that the 40-bp dsRNA provides an interaction surface allowing to simultaneously accommodate more than one VP3 dimer.

A comparison of apparent dissociation constant (K_D_) values recorded with duplexes ranging from 12- to 32-bp (Table 1) indicates that, as predicted by the model, the dsRNA/VP3 binding affinity is largely proportional to duplex length, showing an increase of over 4 log_10_ K_D_ units when comparing data recorded with the 12- and 32-bp dsRNA analytes, respectively.

**Table 1.** Kinetic and affinity constants of the VP3-dsRNA interaction. Presented data were obtained from sensorgrams shown in Supplementary Figure 1 using the BIAevaluation 4.1 software.

| dsRNA length (bp) | $k_a$ ($M^{-1}s^{-1}$) | $k_d$ ($s^{-1}$) | $K_D$ (M) |
| --- | --- | --- | --- |
| 10 | n.d. | n.d. | n.d. |
| 12 | $1.15 \times 10^6$ | $1.76 \times 10^{-1}$ | $1.52 \times 10^{-7}$ |
| 14 | $1.11 \times 10^6$ | $4.48 \times 10^{-2}$ | $4.05 \times 10^{-8}$ |
| 16 | $1.51 \times 10^6$ | $4.50 \times 10^{-2}$ | $3.05 \times 10^{-8}$ |
| 18 | $3.61 \times 10^6$ | $4.41 \times 10^{-2}$ | $1.22 \times 10^{-8}$ |
| 20 | $6.26 \times 10^6$ | $1.32 \times 10^{-2}$ | $2.12 \times 10^{-9}$ |
| 22 | $9.40 \times 10^6$ | $3.02 \times 10^{-2}$ | $3.21 \times 10^{-9}$ |
| 24 | $7.14 \times 10^6$ | $1.42 \times 10^{-2}$ | $2.00 \times 10^{-9}$ |
| 32 | $7.73 \times 10^6$ | $3.60 \times 10^{-3}$ | $3.99 \times 10^{-10}$ |
| 40 | n.d. | n.d. | n.d. |

### Mutational analysis of the VP3 dsRBD

Fitting our previous EMSA data (Valli et al. 2012), the proposed model predicts that both dsRBD subdomains are crucial for the establishment of the dsRNA/VP3 interaction. So, we initiate the mutational assessment of the proposed interaction model by analysing the binding capacity of two previously described mutant VP3 polypeptides, hVP3Patch1 and hVP3Patch2, using SPR (Valli et al. 2012). A cartoon showing the electrostatic three-dimensional surface of wild-type, Patch1 and Patch2, hVP3 dimers used for this analysis is shown in Fig. 2A.

**Figure 2.**
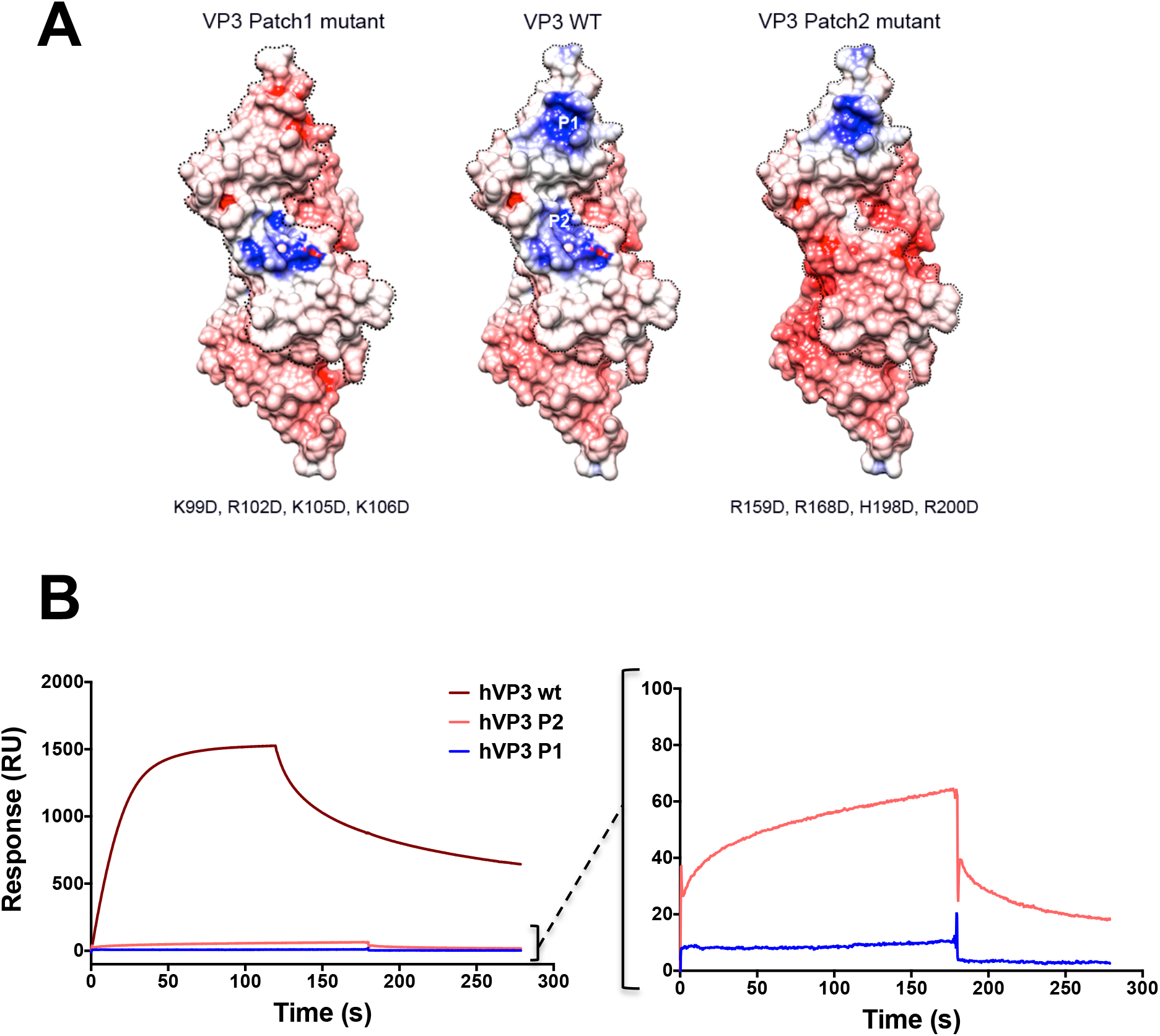
Comparative analysis of the dsRNA binding capacity of Patch1 and Patch2 hVP3 mutant polypeptides. **(A)** hVP3 wt, hVP3Patch1 and hVP3Patch2 protein mutants displayed with its Connolly surface colored according to its electrostatic potential. Residues mutated to alter the electrostatic potential of each hVP3 version are indicated. **(B)** Biacore surface plasmon resonance analysis performed with the wild type (wt), Patch1 (P1) and Patch2 (P2) versions of the hVP3 polypeptide. A biotinylated 24-bp RNA duplex was immobilized on the surface of different cells of a streptavidin sensor-chip. Affinity purified hVP3 protein versions were then injected at 200 nM for 3 min. The graph on the left hand side corresponds to sensograms recorded following the injection of the three proteins under analysis. An enlarged view of sensograms recorded with hVP3Patch1 and hVP3Patch2 polypeptides is presented on the right hand side.

The 24-bp biotinylated dsRNA was captured on the surface of parallel flow cells of a streptavidin sensor-chip as described above. Affinity purified wild type hVP3, hVP3Patch1 and hVP3Patch2 polypeptides were then injected on different flow cells at a concentration of 200 nM for 120 s followed by injection of protein dilution buffer to analyse protein dissociation. An unmodified flow cell served as a reference surface for these experiments. In view of the low resonance signals detected with both hVP3Patch1 and hVP3Patch2 polypeptides, the association phase for these two proteins was extended to 180 s to get a better binding assessment.

Fig. 2B shows a set of representative sensorgrams corresponding to assays performed with the three proteins used in these assays. As expected, a sharp interaction signal was detected with the wild-type hVP3 polypeptide. However, signals gathered with hVP3Patch1 and hVP3Patch2 mutants were exceedingly lower, thus conspicuously revealing the harsh detrimental effect of mutations on both Patch1 and Patch2 dsRBD subdomains. Significantly, the obliteration of Patch1 completely arrests dsRNA binding. In contrast, the hVP3Patch2 mutant protein lacking the Patch2 dsRBD subdomain retains a marginal, yet detectable, capacity to interact with the dsRNA analyte. This observation is consistent with the accessory role of the dsRBD Patch2 subdomain predicted by the interaction model described above.

Next, to assess the specific contribution of individual Patch1 residues, we generated a set of hVP3 polypeptides holding single amino acid substitutions replacing each electropositive residue (K or R) by an electronegative D residue. The resulting protein mutants (hVP3K99D, hVP3R102D, hVP3K105D and hVP3K106D) were tested using parallel flow cells loaded with the 24-bp RNA duplex. Assays were performed by injecting purified proteins at three concentrations (i.e. 50, 100 and 200 nM) using a 60 s association phase. As shown in Fig. 3A, while hVP3R102D and hVP3K105D retain a dose-dependent binding capacity, this activity is completely abolished in their hVP3K99D and hVP3K106D counterparts.

**Figure 3.**
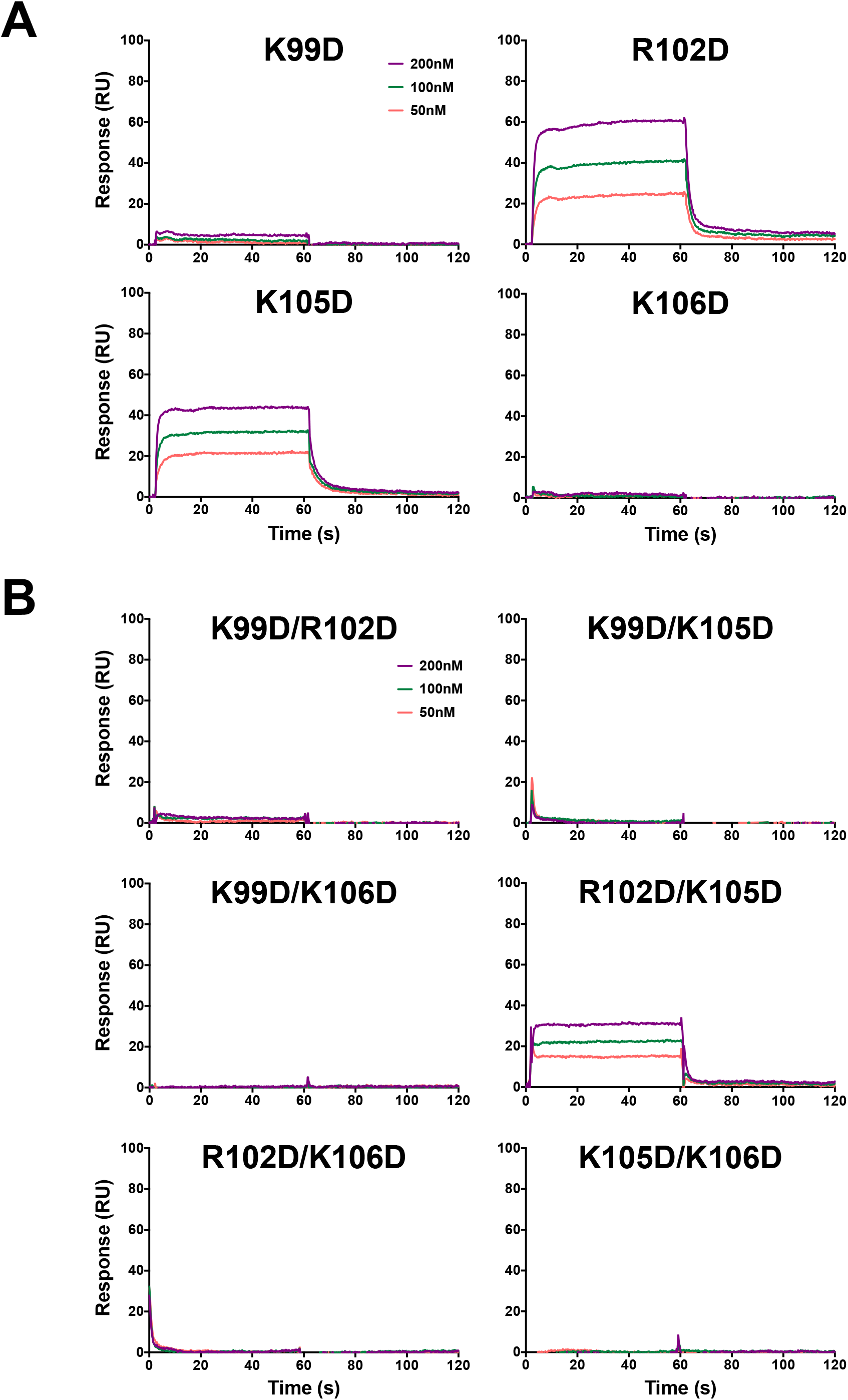
VP3 residues K99 and K106 are necessary for the interaction with the dsRNA. Biacore surface plasmon resonance analysis performed with mutant versions of the hVP3 polypeptide harbouring **(A)** single (hVP3K99D, hVP3R102D, hVP3K105D and hVP3K106D) or **(B)** double (hVP3K99D/R102D, hVP3K99D/K105D, hVP3K99D/K106D, hVP3R102D/K105D, hVP3R102D/K106D and hVP3K105D/K106D) amino acid substitutions on electropositive Patch1 residues. A biotinylated 24-bp RNA duplex was immobilized on the surface of different cells of a streptavidin sensor-chip. Affinity purified hVP3 protein versions were then injected at the indicated concentrations for 1 min.

To further confirm data gathered with single point hVP3 mutants, a set of double mutants within the dsRBD Patch1 subdomain (hVP3K99D/R102D, hVP3K99D/K105D, hVP3K99D/K106D, hVP3R102D/K105D, hVP3R102D/K106D and hVP3K105D/K106D) were generated and tested by SPR. The results of this analysis, shown in Fig. 3B, indicate that the dsRNA binding capacity is exclusively retained by the hVP3R102D/K105D mutant protein. Significantly, the only mutant holding unaltered K99 and K106 residues.

Taken together this set of results show that, as predicted by the VP3-dsRNA interaction model, both K99 and K106 are absolutely essential for VP3-dsRNA complex formation.

### VP3 residues K99 and K106 play a key role counteracting the innate antiviral response

Binding to the viral dsRNA genome represents a key function of VP3 to evade cellular dsRNA sensors that would otherwise trigger the innate antiviral response. In light of the different dsRNA binding activities observed for the tested VP3 mutant versions, it seemed feasible to hypothesize a correlation between the dsRNA-binding affinity and the capacity to control the activation of the innate antiviral response. Accordingly, we compared the capacity of VP3 and its derived mutant versions to hamper the activation of the innate antiviral activity triggered by dsRNA transfection (Fig. 4). For this, we designed an experimental approach allowing to quantify the transcriptional activation of the IFN-β promoter in response to synthetic Poly I:C dsRNA transfection.

**Figure 4.**
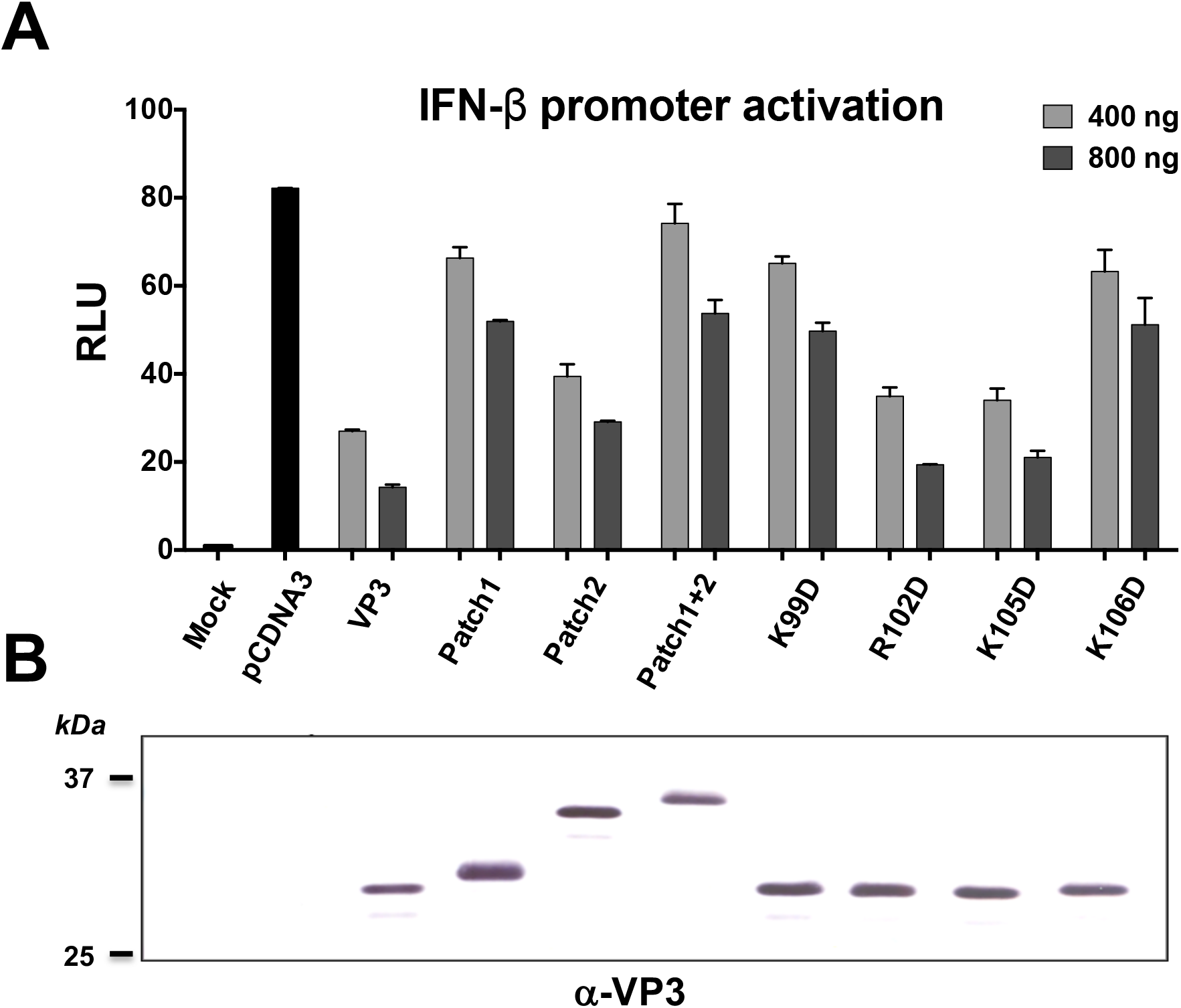
VP3 residues K99 and K106 are essential to counteract the innate antiviral response. (A) IFN-ß promoter activation. DF-1 cells were transfected with a plasmid expressing the *Photinus pyralis* luciferase gene under the control of the human IFN-β promoter, along with the pRL-null plasmid, constitutively expressing the *Renilla muelleri* luciferase gene. The latter used as transfection control. Cells were simultaneously transfected with 400 or 800 ng of pCDNA3 derived plasmids expressing the different VP3 protein versions, or with identical amounts of empty pcDNA3. At 8 h post-transfection, IFN-β promoter activity was induced by transfecting 250 ng of poly I:C. After 16 h, cell lysates were collected, and luciferase activities were recorded. *Photinus* luciferase activity was expressed as the relative fold induction (n-fold) over that detected in the pcDNA3-transfected cells, after normalization to the *Renilla* luciferase activity. Charts correspond to the mean ±the standard deviation of three independent experiments. **(B) Expression of VP3 variants.** Western blot analysis (α-VP3) of the expression of VP3wt and indicated mutants in DF-1 cells.

DF-1 chicken cells were cotransfected with pIFNβ-luc, expressing the *Photinus pyralis* luciferase gene (PL) under the control of the IFN-β promoter, and a pcDNA3-derivative constitutively expressing each VP3 version under analysis; pRL-null, constitutively expressing the *Renilla muelleri* (RL) luciferase gene was used as transfection control. 18 h later, cultures were transfected with Poly I:C to trigger the transcriptional activation of the IFN-β promoter. After 16 h, cells were collected and the corresponding extracts used to quantify both PL and PR activities.

As shown in Fig. 4, transfection of poly I:C induced a strong activation of the IFN-β promoter in cells transfected with the empty pcDNA3 vector control. This activation is strongly reduced in the presence of wild type VP3, suggesting that the synthetic dsRNA is sheltered by the protein and thus, not detected by cellular sensors. As expected, expression of the VP3Patch1 mutant, unable to bind dsRNA, shows high levels of IFN-β promoter activation, almost as high as those found with the empty vector control; and in the case of the VP3Patch2 mutant, IFN-β activation levels are significantly reduced.

Interestingly, when expressing the VP3 versions bearing single amino acid substitutions on the Patch1 dsRBD subdomain we observed that only VP3R102D and VP3K105D were able to reduce IFN-β promoter activation to levels similar to those achieved with the VP3 wild type. VP3K99D and VP3K106D mutants behaved similarly to VP3Patch1, and in both cases poly I:C transfection induced a strong activation of the IFN-β promoter due to the sheltering deficiency of both mutant proteins.

In all cases, a VP3 dose-dependent response is observed, further supporting the correlation between the capacity to interact with dsRNA and the evasion from the cellular sensors. Remarkably, the substitution of only one residue, either K99 or K106, completely abolishes the capacity of the protein to evade the antiviral response. Indeed, this finding offers a new and interesting therapeutic target that could be further explored in the future.

### VP3 residues K99 and K106 are essential for IBDV replication

In view of results described above it was of outmost importance to determine the effect that K99D and K106D mutations might exert on IBDV replication. For this, we used a previously described reverse genetics approach based on the use of two plasmids, i.e. pT7_SA_Rz and B pT7_SB_Rz, harbouring cDNA sequences corresponding to IBDV genome segments A and B under the transcriptional control of the T7 bacteriophage RNA polymerase (Irigoyen et al. 2009). VP3K99D and VP3K106D substitutions were introduced into the IBDV segment A coding sequence by site-directed mutagenesis. The resulting plasmids pT7_SA-VP3K99D_Rz and pT7_SA-VP3K106D_Rz respectively, were used to cotransfect QM7 cells along with the plasmid pT7_SB_Rz, expressing segment B. As a control for these experiments, cultures were cotransfected with plasmids pT7_SA_Rz and pT7_SB_Rz expressing cDNAs corresponding to wild-type IBDV genome segments A and B, respectively. After transfection, cultures were infected with VT7LacOI, a recombinant vaccinia virus expressing the T7 RNA polymerase upon addition of IPTG to cell media. Cultures were then maintained in the presence of the IPTG to trigger transcription of both IBDV genome segments. At 72 h PI, cell supernatants were harvested and used to detect infectious IBDV by plaque assay. Supernatants were also used to further amplify infectious IBDV by two consecutive rounds of infection on fresh QM7 cell cultures. Results presented in Table 2, indicate that both, VP3K99D and VP3K106D, mutations are lethal, completely abolishing the production of an infective IBDV progeny.

**Table 2.** Reverse genetics analysis of the effect of VP3K99D and VP3K106D mutations on the rescue of infectious IBDV. QM7 cells were transfected with a combination of plasmids pT7-SA-Rz and pT7-SB-Rz, pT7-SAVP3K99D-Rz and pT7-SB-Rz, or pT7-SAVP3K106D-Rz and pT7-SB-Rz, expressing VP3 wild-type, VP3K99D or VP3K106D respectively. At 6 h post-transfection, cultures were infected with 3 PFU/cell of the rVACV VT7. At 72 h PI, supernatants were collected and used to infect fresh QM7 cell monolayers. Samples from these infections were used to perform two subsequent rounds of IBDV amplification by infecting fresh QM7 monolayers. The presence of IBDV was assessed by plaque assay. Presented data corresponds to virus titrations performed in triplicate from samples collected from three independent experiments. (N.D.: Not Detected)

| VP3 gene | Virus titers (PFU/ml) |  |  |
| --- | --- | --- | --- |
|  | First round | Second round | Third round |
| Wild-type | $2.1 \pm 1.2 \times 10^2$ | $3.4 \pm 2.3 \times 10^4$ | $6 \pm 3.1 \times 10^7$ |
| VP3K99D | N.D. | N.D. | N.D. |
| VP3K106D | N.D. | N.D. | N.D. |

Western blotting analysis performed with samples from transfected/infected cultures showed that the mutations under analysis do not affect VP3 expression. This rules out the possibility that the failure to recovering infectious IBDV might be due defects on the expression of mutant VP3 genes (Fig. 5A).

**Figure 5.**
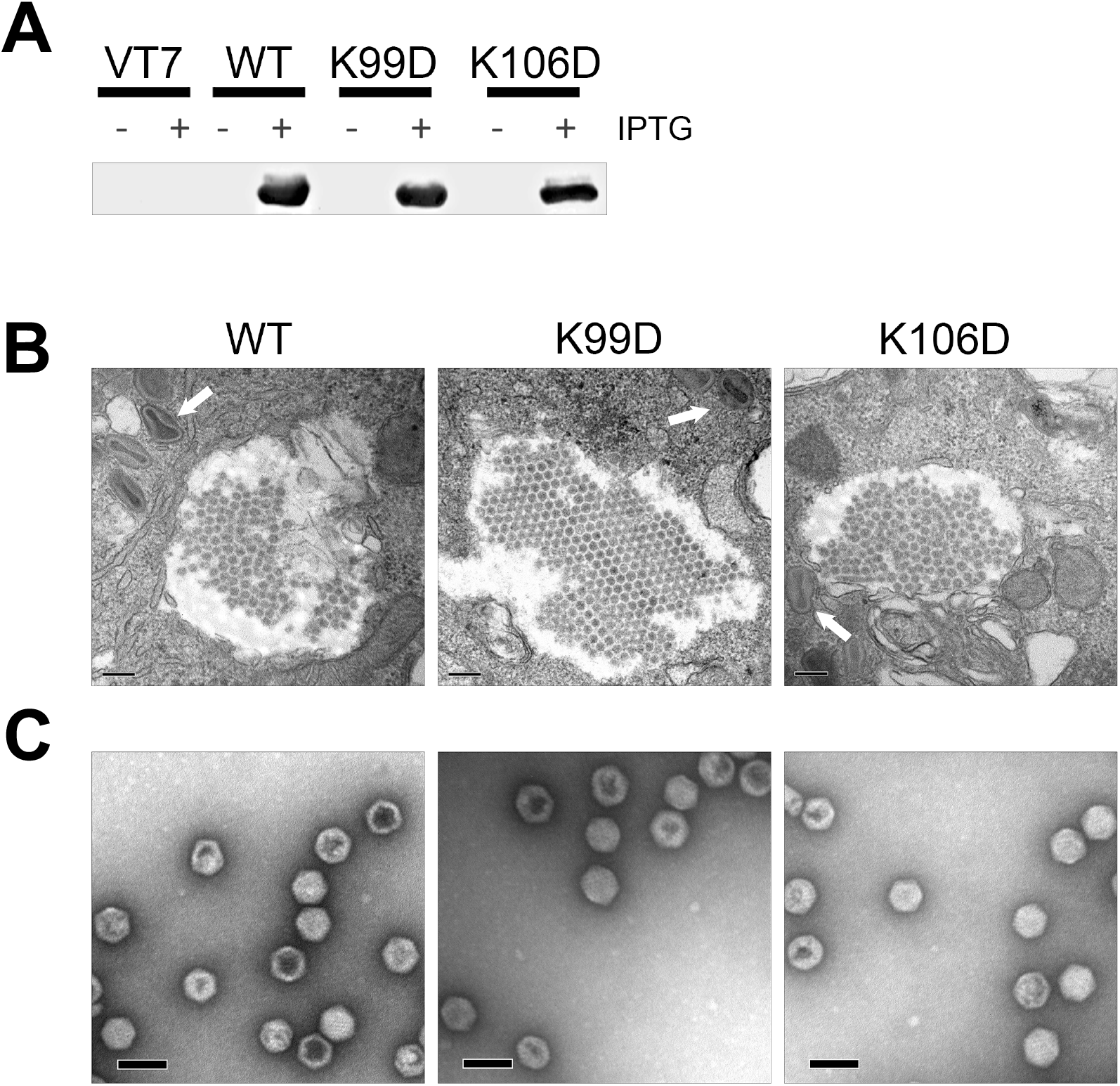
K99D or K106D mutations in VP3 do not alter its scaffolding activity during IBDV particle morphogenesis. **(A)** Western blot analysis performed with the anti-VP3 serum using lysates from QM7 cells co-transfected with plasmids pT7_SA_Rz and pT7_SB_Rz (WT); or pT7_SA-VP3K99D_Rz and pT7_SB_Rz (K99D); or pT7_SA-VP3K106D and pT7_SB_Rz (K106D) and infected with VT7LacOI in the presence (+) or absence (-) of IPTG. Untransfected VT7LacOI-infected cells (VT7) were used as control. **(B)** BSC-1 cell monolayers were infected with VT7_POLY (WT), VT7_POLY-VP3K99D (K99D) or VT7_POLY-VP3K106D (K106D) and maintained for 48 h in the presence of IPTG. Cultures were then processed for transmission electron microscopy. Panels show ultrathin sections corresponding to the cytoplasm of cells infected with the different viruses. Arrows indicate the position of vaccinia virus virions. Scale bars represent 200 nm. **(C)** Electron microscopy images of negatively stained IBDV-like particles purified from BSC-1 cultures infected with VT7_POLY (WT), VT7_POLY-VP3K99D (K99D) or VT7_POLY-VP3K106D (K106D). Scale bars represent 50 nm.

The VP3 polypeptide acts as an essential scaffolding element during IBDV particle assembly (Oña et al. 2004). Accordingly, it seemed feasible that mutations under analysis might alter capsid morphogenesis. Therefore, we next analysed the effect of both VP3K99D and VP3K106D on the assembly of IBDV virus-like particles (VLP). Two recombinant vaccinia viruses VT7_POLY-VP3K99D and VT7_POLY-VP3K106D, expressing mutant versions of the IBDV polyprotein gene, harbouring either the VP3K99D or the VP3K106D point mutations, were generated. The previously described VT7_POLY recombinant vaccinia virus, expressing the wild-type polyprotein gene, known to direct the assembly of IBDV VLP (Fernández-Arias et al. 1998), was used as a control for subsequent experiments. These three recombinant viruses were used to infect cultures of BSC-1 cells. Infected cultures were maintained in the presence of the IPTG to trigger the expression of the different polyprotein gene versions.

IBDV capsid assembly was tested using two complementary approaches: i) transmission electron microscopy analysis of cells infected with the described recombinant vaccinia viruses; and ii) sucrose gradient-based purification of VLP from infected cell extracts. As shown in Fig. 5B, regardless of the polyprotein gene version being expressed, typical honeycomb-like IBDV-derived VLP superstructures were detected within the cytoplasm of infected cells. Additionally, icosahedral VLPs identical to *bona fide* IBDV particles were isolated from extracts corresponding to cells expressing all three genes under analysis (Fig. 5C). Hence, showing that neither K99D nor K106D affect the VP3 scaffolding activity.

## DISCUSSION

RNPs are a distinctive structural and functional hallmark differentiating Birnaviruses from prototypical dsRNA viruses deserving an in depth characterization. Indeed, the lack of VP3-dsRNA complex model posed a major obstacle to both understanding mechanisms governing RNP assembly/disassembly processes and to exploring the RNP’s contribution to the birnavirus replication process.

Here, we present a hypothetical model based on the *in silico* docking of the VP3 dimer on the A-form dsRNA helix suggesting that complex formation relies on the establishment of electrostatic interactions between the RNA duplex and VP3 dimers. The first one involving OH^-^ groups from 5’-phosphates of nucleotides n1 and n9c, within the major dsRNA groove, and electropositive side chains of VP3 residues K99 and K106, located at the Patch1 region. According to the model, a second interaction is established between the OH^-^ group from 5’-phosphate of n17c, also placed at the RNA’s major groove, and electropositive Patch2 residues. In this scenario, the interaction mediated by the dsRBD Patch1 subdomain would play a major role, docking the VP3 dimer on the major groove of the RNA helix, whilst the Patch2 subdomain would have a secondary function stabilizing the complex by further engaging the protein dimer on the major groove of the next helix turn. The model predicts that complex assembly to be strictly dependent on the length of the RNA duplex. So, stable complexes, engaging both VP3 Patch1 and 2 regions, would only be formed with dsRNA molecules offering a perfectly base-paired helix comprising at least 17-bp. VP3 dimers should also weakly interact with smaller RNA duplexes holding a flawless helical structure comprising at least 9-bp.

The robustness of the proposed model was tested using a SPR-based approach using both a range of dsRNA analytes of different sizes and recombinant VP3 polypeptides harbouring mutations on the previously described dsRNA subdomains.

Results gathered with the different dsRNA analytes are conclusive showing that, as predicted by the model, VP3/dsRNA complex formation is utterly dependent on dsRNA length. Although predictions concerning helix size requirements are bound to be affected by the inherent dsRNA instability, experimental data roughly fit the proposed model. Hence, showing a clear correlation between the size of the dsRNA analytes and the VP3 binding affinity. The analysis performed with VP3 mutant protein versions confirm the involvement of both dsRBD subdomain, Additionally, as predicted by the model, the analysis show that the electropositive Patch1 subdomain plays a chief role in complex formation. Data collected from the analysis of mutant VP3 polypeptides harbouring Patch1 single and double amino acid substitutions demonstrate that, as predicted by the *in silico* model, residues K99 and K106 are absolutely essential for the interaction to take place. Taking together SPR data presented are in good agreement with the proposed dsRNA/VP3 interaction model.

### The role of RNPs on IBDV replication

As described above, experimental data gathered during the assessment of the dsRNA/VP3 interaction model showed the essential role of both VP3 residues K99 and K106. Indeed, the replacement of either one of these residues by an electronegative amino acid completely abrogates the dsRNA binding capacity of the protein. We took advantage of this finding in order to analyse the effect of such mutations on the capacity of the protein to hinder the recognition of dsRNA by cellular dsRNA sensors and the subsequent activation of the innate antiviral response. The results of this analysis conclusively show that both residues are critical to counteracting the dsRNA-triggered innate host’s response. Moreover, here we show that although these mutations do not affect the VP3 scaffolding activity, allowing capsid morphogenesis, their introduction within the virus genome completely preclude virus replication. Hence showing for the first time how the introduction of a single amino acid substitution (either K99D or K106D) efficiently abrogates the ability of the virus to complete its replication programme. Certainly, these two critical amino acid residues offer a highly promising target for the design of structure-based of IBDV-specific antivirals. Indeed, data presented here underscore the utmost importance of RNP complexes during the IBDV life cycle.

### Insights into the IBDV replication cycle

According to SPR data, VP3-dsRNA association and dissociation phases are fast and highly dependent upon protein concentration. This was somehow expected due to the predicted roles of RNPs during the IBDV replication process; i.e. acting as transcription/replication devices, hindering dsRNA sentinel cellular proteins and likely cooperating to the encapsidation of the virus genome.

In order to facilitate the transcription/replication process, VP3 dimers should readily dislodge from dsRNA, thus allowing the synthesis of the nascent RNA daughter chain without affecting the processivity of the viral RdRp. The apparent K_D_ of VP3 dimer interaction with duplexes ranging from 20- to 24-bp dsRNA is ca. 2 nM. Similar studies conducted with other virus-encoded RNA-interacting proteins rendered kinetic constants comparable to that detected with VP3, e.g. the structural NP protein of Influenza virus (K_D_= 8.27 ± 1.43 nM) (Liu et al. 2016); or the N nucleocapsid protein of coronaviruses (K_D_= 0.66-2.82 nM) (Chen et al. 2005; Spencer and Hiscox 2006). Although experimental differences preclude a direct comparison, our data indicate that the affinity VP3 for dsRNA is not too far off from those recorded with other viral RNA-binding proteins.

Additionally, VP3 dimers should swiftly bind the newly synthesized dsRNAs, thus preventing dsRNA detection. Indeed, VP3 dimer binding should be favoured by the high concentration of this protein detected at IBDV replication sites (Dalton and Rodríguez 2014).

The proposed VP3-dsRNA interaction model suggests that IBDV replication leads to formation of a complex network built by intertwined genome segments, reversibly glued by VP3 dimers within the cell cytoplasm. Such network would act as a molecular trap engulfing newly synthesized RNP components, i.e. dsRNA, VP3 and VP1. This probably explains the presence of rather large viral factories within discrete cytoplasmic areas of IBDV infected cells (Méndez et al. 2017). In addition to conferring an effective shield against specialized host’s dsRNA sensors, this RNP network might also serve as a convenient platform to facilitate interactions between pVP2, the capsid precursor polypeptide, and RNP-associated VP3 dimers, probably facilitating the previously described random packaging of genome segments (Luque et al. 2009) during the IBDV assembly process.

According to previously published data, the large majority of infectious IBDV particles package four genome segments in a random manner (Luque et al. 2009). This accounts for a total RNA duplex length of ca. 12,200-bp per particle (four segments with an average length of 3,050-bp). We have also shown that IBDV virions contain 457±50 VP3 monomers (228±25 dimers) (Luque et al. 2009). Each VP3 dimer possesses two dsRBDs, one at each dimer face, thus being able to simultaneously interact with two dsRNA molecules. Our experimental data indicates that the RNA duplex size stably holding two VP3 dimers is of 20-24-bp. Provided this is maintained under natural infection conditions, a simple calculation, based on available genome and VP3 stoichiometric data, indicates that the number of VP3 dimers required to completely enfold the four dsRNA genome segments found in IBDV particles should be between 305 and 254, quite close to the figure found in purified IBDV virions. This suggests that RNPs released into the cytoplasm from infecting virus particles are fully protected by VP3 dimers. Hence, providing a mechanism to ensuring the stealth of the initial virus replication steps.

## MATERIALS AND METHODS

### Cells and viruses

BSC-1 (African green monkey kidney cells, ATCC number CRL-2761), DF-1 (spontaneously transformed chicken embryo fibroblasts, ATCC number CRL-12203) and QM7 (quail muscle myoblasts, ATCC number CRL-19DF cells were grown in Dulbecco’s modified minimal essential medium (DMEM). *HighFive* cells (*Trichoplusia ni* ovary cells, BTI-TN-5B1-4, Invitrogen) were grown in TC-100 medium (GIBCO). All cell media were supplemented with penicillin (100 U/ml), streptomycin (100 μg/ml) and 10% fetal calf serum (FCS) (Sigma). The recombinant VACV VT7LacOI, kindly provided by B. Moss (National Institute of Health, Bethesda, Maryland, USA), was grown and titrated in BSC-1 cells as previously described (Ward et al. 1995). All recombinant baculoviruses (rBV) used in this report were grown and titrated in HighFive cells following instructions provided in the Bac-to-Bac Baculovirus Expression System manual (Invitrogen, Publication Number MAN0000414. Revision A.0.). IBDV infections and titrations were carried out in QM7 cells as previously described (Méndez et al. 2015).

### Generation of rBVs

rBVs FB/his-VP3, FB/his-VP3P1 and FB/his-VP3P2, used for the production of recombinant hVP3, hVP3P1 and hVP3P2 have been described elsewhere (Valli et al. 2012; Kochan et al. 2003). Other rBVs used in this report were generated as follows. The construction of baculovirus transfer vectors encoding hVP3 mutant polypeptides were generated using PCR-based site directed mutagenesis on the pFB/hisVP3 plasmid (Kochan et al. 2003) using synthetic DNA oligonucleotide primers (Sigma) described in Supplementary Table 1. The resulting plasmids, pFB/hVP3K99D, pFB/hisVP3R102D, pFB/hVP3K105D and pFB/hVP3K106D, respectively, were used for the introduction of double mutations using the corresponding oligonucleotide primers (Supplementary Table 1), thus generating plasmids pFB/hVP3K99D-R102D, pFB/hVP3K99D-K105D, pFB/hVP3K99D-K106D, pFB/hVP3R102D-K105D, pFB/hVP3R102D-K106D and pFB/hisK105D-K106D). Plasmids were subjected to nucleotide sequencing and then used to generate rBVs using the Bac-to-Bac system (Invitrogen).

### Generation of pcDNA3-VP3 plasmids

Generation of pcDNA3-VP3wt has been described elsewhere (Busnadiego et al. 2012). For the generation of pcDNA3VP3Patch1, pcDNA3VP3Patch2, pcDNA3VP3Patch1+2, pcDNAVP3K99D, pcDNAVP3R102D, pcDNAVP3K105D and pcDNAVP3K106D, DNA fragments corresponding to each of the VP3 coding regions were generated by PCR from the previously described pFB baculovirus transfer vectors using the primers 5’-CGCGAAGCTTATGGGTTTCCCTCACAATCCACGC and 5’-GCGCGGATCCTCACTCAAGGTCCTCATCAGAGAC. The DNA fragments were purified, restricted with HindIII and BamHI and cloned into pcDNA3 (Invitrogen) previously digested with the same enzymes. The resulting plasmids were subjected to nucleotide sequence analysis to assess the correctness of the cloned sequence.

### Generation of recombinant VACV expressing mutant versions of the VP3 polypeptide

The production of recombinant VACV (rVACV) VT7/VP3K99D and VT7/VP3K106D, was initiated by generating the corresponding VACV insertion vectors. For this, two DNA fragments of 789 bp containing the VP3K99D and VP3K106D mutant versions of the VP3 ORF, flanked by NdeI and BamHI restriction sites, were generated by PCR using the plasmids pFB/hisVP3K99D and pFB/hisVP3K106D as templates and the primers 5’GCGCCATATGGCTGCATCAGAGTTCAAAGAG and 5’GCGCGGATCCTCACTCAAGGTCCTCATCAGAG. The resulting PCR products were purified, digested with NdeI and BamHI and ligated to the pVOTE.2 VACV insertion/expression plasmid vector (Ward et al. 1995) previously digested with the same restriction enzymes. The resulting plasmids, pVOTE/VP3K99D and pVOTE/VP3K106D, were subjected to nucleotide sequencing, and then used to generate the corresponding rVACVs, VT7/VP3K99D and VT7/VP3K106D, via homologous recombination with the VT7 VACV genome as previously described (Ward et al. 1995).

### Expression and Purification of his-tagged VP3 protein versions from Insect cells

Expression of the different hVP3 protein versions was achieved by infecting HighFive cell monolayers with the rBVs described above at a MOI of 3 PFU/cell. Infected cells were harvested at 72 h post-infection (PI), washed twice with phosphate-buffered saline, resuspended in lysis buffer (50 mM Tris-HCl [pH 8.0], 500 mM NaCl, 0.1% igepal) supplemented with protease inhibitors (Complete Mini; Roche), and maintained on ice for 20 min. Thereafter, extracts were centrifuged at 13,000xg for 10 min at 4°C. Supernatants were collected and subjected to metal-affinity chromatography (IMAC) purification by using a Ni^2+^ affinity column (HisTrap HP, GE Healthcare). Resin-bound polypeptides were released with elution buffer (50 mM Tris-HCl [pH 8.0], 500 mM NaCl, 250 mM imidazol). hisVP3-containing fractions were pooled, dialyzed against lysis buffer lacking igepal, and subjected to a second purification round under identical conditions. Finally, protein samples were dialyzed against 1,000 volumes of 50 mM Tris-HCl (pH 8.0), 150 mM NaCl. The purity of eluted proteins was assessed by SDS-PAGE analysis and Coomassie blue staining. Protein concentration was determined using the BCA system (Thermo Scientific Pierce, USA). Purified proteins were maintained at 4°C for a maximum period of 14 days.

### Generation of dsRNA analytes

RNA duplex preparation was performed by annealing complementary synthetic single stranded (ss) RNA oligonucleotide pairs where one of the oligonucleotides from each pair was biotinylated at its 5’-end. Both oligonucleotide synthesis and annealing were performed by the manufacturer (Biomers.net). Oligonucleotide sequences are described in Supplementary Table 2.

### Surface Plasmon Resonance Analysis

SPR experiments were performed using a biosensor Biacore 3000 (Biacore, GE Healthcare). dsRNA ligands bearing a biotin at the 5’end of one the strands were immobilized on the surface of streptavidin-coated sensor chips (SA). RNA duplexes were loaded at a flow rate of 10 μl/min, using HBS-P (10 mM HEPES [pH 7.4], 0.2 M NaCl, 3 mM EDTA, 0.005% Surfactant P20) running buffer. Reference surfaces, used as control for these assays, were blank flow cells.

Binding assays, performed in duplicate within each experiment, were carried out at 25°C at a flow rate of 30 μl/min. Proteins under study were serially diluted in running buffer to reach the indicated concentrations. The protein fraction that remained bound after the dissociation phase was removed by injecting 800 mM NaCl. Data were collected for the association and dissociation periods as indicated in each specific experiment. SPR sensograms recorded with each tested protein concentration were overlaid, aligned and analysed using the BIAevaluation Software 4.1 (GE Healthcare). All data sets were processed using a double-referencing method (Myszka 2000). Collected data were fit to a 1:1 Langmuir model with a correction for mass transport (Morton et al. 1995).

### IBDV reverse genetics analysis

The reverse genetics analysis was performed as previously described (Irigoyen et al. 2009) using an approach based on the co-transfection of QM7 cells with two plasmids containing cDNAs corresponding to the positive strand sequence of the IBDV segments A and B, respectively. Plasmid pT7-SA-Rz harboring the cDNA of segment A was subjected to site directed mutagenesis to introduce mutations VP3K99D or VP3K106D within the context of the IBDV polyprotein open reading frame. Mutagenesis was performed as described above using primer pairs K99DFW and K99DREV or K106DFW and K106DREV (Supplementary Table 1). The resulting plasmids, pT7-SAVP3K99D-Rz and pT7-SAVP3K106D-Rz, were subjected to nucleotide sequence analysis to assess their correctness.

QM7 cells were transfected with a combination of plasmids pT7-SA-Rz and pT7-SB-Rz, corresponding to the IBDV wild-type genome segments, pT7-SAVP3K99D-Rz and pT7-SB-Rz, or pT7-SAVP3K106D-Rz and pT7-SB-Rz, respectively, using Lipofectamine 2000 (Invitrogen). At 6 h post-transfection, cultures were infected with 3 PFU/cell of VT7, an rVV inducibly expressing the T7 RNA polymerase. Cultures were then maintained at 37°C in medium supplemented with 1 mM isopropyl-D-thiogalactosidase (IPTG). At 72 h PI, cultures were harvested and subjected to three freeze-thaw cycles. Infected cell samples were used to assess the expression of the VP3 polypeptide by Western blotting. After removing cell debris by low speed centrifugation, supernatants were recovered and filtered through 0.1 μm filters (Merk Millipore) to eliminate contaminant rVV particles, and used to infect fresh QM7 cell monolayers. Samples from these infections were collected at 72 h PI and used to rule out possible contamination with the rVACV by Western blotting by using the monoclonal antibody mAbC3 recognizing a highly abundant VACV structural protein (Rodríguez et al. 1985). The corresponding cell supernatants were used to perform two subsequent rounds of IBDV amplification by infecting fresh QM7 monolayers. The presence of IBDV in all samples was assessed by plaque assay as previously described (Méndez et al. 2015).

### Characterization of IBDV virus-like particles in cells infected with rVACV

BSC-1 cell monolayers were infected with rVACV VT7/VP3, VT7/VP3K99D or VT7/VP3K106D at a MOI of 3 PFU/cell. Transmission electron microscopy (TEM) analysis was performed using cell samples collected at 48 h PI following a previously described protocol (Oña et al. 2004). Purification of IBDV virus-like particles (VLP) was carried out using samples collected at 72 h PI as previously described (Fernández-Arias et al. 1998). VLP samples were negatively stained with 2% aqueous uranyl acetate. TEM micrographs were recorded with a Jeol 1200 EXII electron microscope operating at 100 kV.

### Western Blot analysis

Samples used for Western blot analysis were prepared as described previously (Busnadiego et al. 2012). The antibody used in this study was a rabbit polyclonal serum specific for IBDV VP3 (Fernández-Arias et al. 1998). After incubation with the primary antibody, membranes were incubated with goat anti-rabbit IgG-Peroxidase conjugate (Sigma). Immunoreactive bands were detected by enhanced chemiluminescence (GE Healthcare).

### IFN-β promoter activation assay

DF-1 cells were transfected with 400 or 800 ng of plasmid pIF(−116/+72)lucter (pIFNβ-luc), expressing the *Photinus pyralis* luciferase gene under the control of the human IFN-β promoter (King and Goodbourn 1992), along with 30 ng of pRL-null, constitutively expressing the *Renilla muelleri* luciferase gene (Promega). The latter used as transfection control. Cells were simultaneously transfected with 400 or 800 ng of pCDNA3 derived plasmids expressing the different VP3 protein versions, or with identical amounts of empty pcDNA3. All samples received the same amount of total DNA, that was adjusted by adding the required amount of pcDNA3. At 8 h posttransfection, IFN-β promoter activity was induced by transfecting 250 ng of poly I:C (average size 0.2-1 kb) (InvivoGen) during 16 h. After this period, cell lysates were collected and analysed using the dual-luciferase assay kit (Promega) following the manufacter’s instructions. Luciferase activities were recorded using an Orion II luminometer (Titerthek Berthold). *Photinus* luciferase activity was expressed as the relative fold induction (n-fold) over that detected in the pcDNA3-transfected cells, after normalization to the *Renilla* luciferase activity.

## ACKNOWLEDGEMENTS

We are grateful to the excellent technical assistance provided by Antonio Varas.

