## Supplementary material for "Modelling the interaction between Infectious Bursal Disease Virus VP3 polypeptide and dsRNA: identification of key residues for ribonucleoprotein assembly and virus replication": Suppemental Data.pdf (from another version of this preprint: version 1, 2020-08-07)

**Supplemental Figure 1. Effect of RNA duplex length on VP3-dsRNA complex formation.** Biacore surface plasmon resonance analysis of the interaction between the hVP3 polypeptide with RNA duplexes of lengths ranging from 12- to 40-bp. Sensograms were recorded following the injection of affinity purified hVP3 protein at concentration ranging from 20 nM to 0.625 nM, as indicated at the upper right side of the panel, on sensor chips loaded with the different RNA duplexes.

**Supplemental Table 1.** Synthetic DNA oligonucleotides used for the generation of mutant hVP3 polypeptide versions.

**Supplemental Table 2.** Synthetic RNA oligonucleotide pairs used for Biacore surface plasmon resonance analysis.

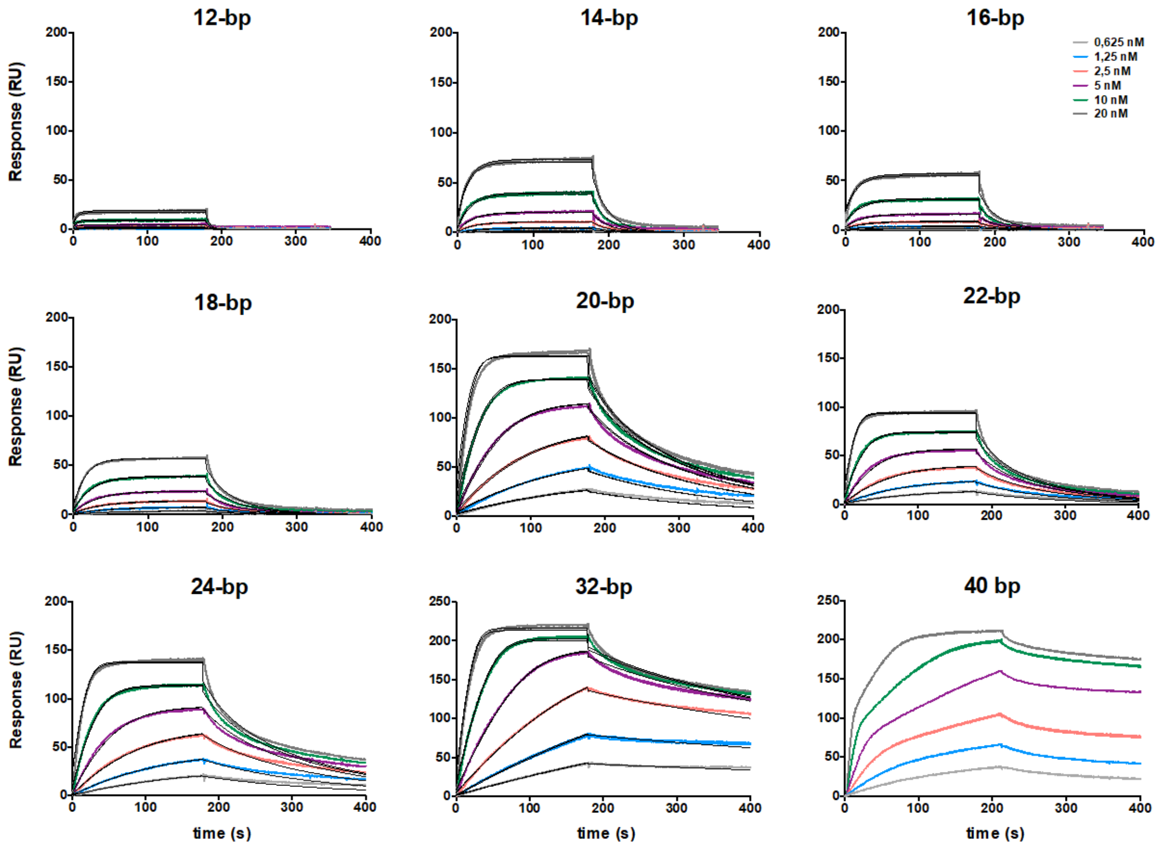


Supplemental Figure 1

| Name | Sequence 5´-3´ | Mutation |
|---|---|---|
| K99D FW | GAAGCACAGAGGGAAGACGACACACGGATCTCA | K99D |
| K99D REV | TGAGATCCGTGTGTCGTCTTCCCTCTGTGCTTC | |
| R102D FW | AGGGAAAAAGACACAGACATCTCAAAGAAGATG | R102D |
| R102D REV | CATCTTCTTTGAGATGTCTGTGTCTTTTTCCCT | |
| K105D FW | GACACACGGATCTCAGACAAGATGGAGACCATG | K105D |
| K105D REV | CATGGTCTCCATCTTGTCTGAGATCCGTGTGTC | |
| K106D FW | ACACGGATCTCAAAGGACATGGAGACCATGGGC | K106D |
| K106D REV | GCCCATGGTCTCCATGTCCTTTGAGATCCGTGT | |
| K99D/R102D FW | AGGGAAGACGACACAGACATCTCAAAGAAGATG | K99D/R102D |
| K99D/R102D REV | CATCTTCTTTGAGATGTCTGTGTCGTCTTCCCT | |
| K99D/K105D FW | GACACACGGATCTCAGACAAGATGGAGACCATG | K99D/K105D |
| K99D/K105D REV | CATGGTCTCCATCTTGTCTGAGATCCGTGTGTC | |
| K99D/K106D FW | ACACGGATCTCAAAGGACATGGAGACCATGGGC | K99D/K106D |
| K99D/K106D REV | GCCCATGGTCTCCATGTCCTTTGAGATCCGTGT | |
| R102D/K105D FW | GACACAGACATCTCAGACAAGATGGAGACCATG | R102D/K105D |
| R102D/K105D REV | CATGGTCTCCATCTTGTCTGAGATGTCTGTGTC | |
| R102D/K106D FW | ACAGACATCTCAAAGGACATGGAGACCATGGGC | R102D/K106D |
| R102D/K106D REV | GCCCATGGTCTCCATGTCCTTTGAGATGTCTGT | |
| K105D/K106D FW | ACACGGATCTCAGACGACATGGAGACCATGGGC | K105D/K106D |
| K105D/K106D REV | GCCCATGGTCTCCATGTCGTCTGAGATCCGTGT | |

**Supplemental Table 1.** Synthetic DNA oligonucleotide pairs used for the introduction of mutations within the hVP3 gene.

Supplemental Table 1

| Name | Sequence 5´-3´ |
|---|---|
| 10B | UCACUUACGC |
| 10RC | GCGUAAGUGA |
| 12B | UCACUUACGCUG |
| 12RC | CAGCGUAAGUGA |
| 14B | UCACUUACGCUGAG |
| 14RC | CUCAGCGUAAGUGA |
| 16B | UCACUUACGCUGAGUA |
| 16RC | UACUCAGCGUAAGUGA |
| 18B | UCACUUACGCUGAGUACU |
| 18RC | AGUACUCAGCGUAAGUGA |
| 20B | UCACUUACGCUGAGUACUUC |
| 20RC | GAAGUACUCAGCGUAAGUGA |
| 22B | UCACUUACGCUGAGUACUUCGA |
| 22RC | UCGAAGUACUCAGCGUAAGUGA |
| 24B | UCACUUACGCUGAGUACUUCGAGA |
| 24RC | UCUCGAAGUACUCAGCGUAAGUGA |
| 32B | UCACUUACGCUGAGUACUUCGAGACUUCGAGA |
| 32RC | UCUCGAAGUCUCGAAGUACUCAGCGUAAGUGA |
| 40B | UCACUUACGCUGAGUACUUCGAGACUUCGAGACUUCGAGA |
| 40RC | UCUCGAAGUCUCGAAGUCUCGAAGUACUCAGCGUAAGUGA |

**Supplemental Table 2.** Synthetic RNA oligonucleotide pairs used for the generation of RNA duplexes. B indicates biotinylated oligonucleotides

Supplemental Table 2
